## Supplementary material for "Fitting a lattice model with local and global transmission to spread of a plant disease"

### Supplementary Plots

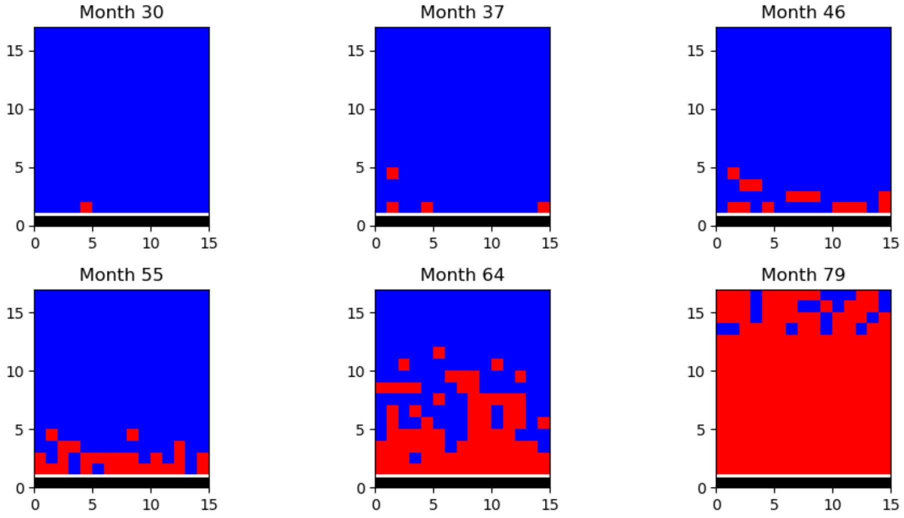


Figure 1: **S1: Visualisation of experimental data.** Red squares denote symptomatic trees and blue squares non-symptomatic, with the row of founder trees in black. Note that unlike the model snapshots it is not possible to detect trees in their latent infected period.

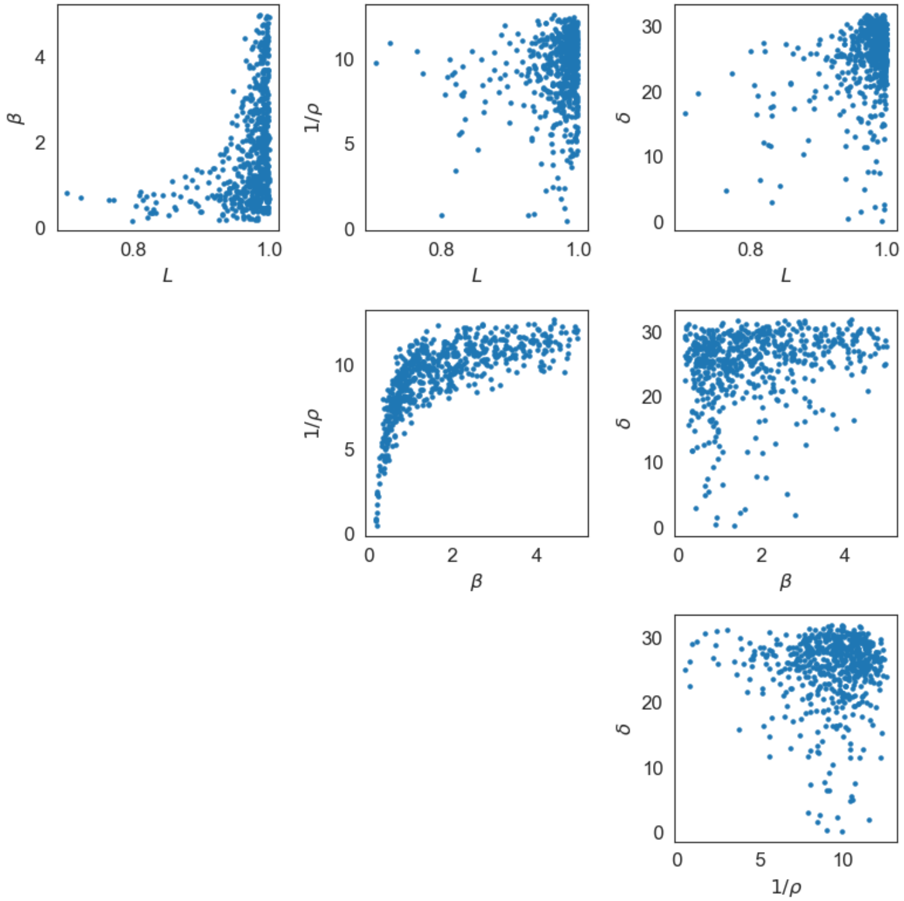


Figure 2: **S2: Correlations between the posterior parameters from the lattice model.**

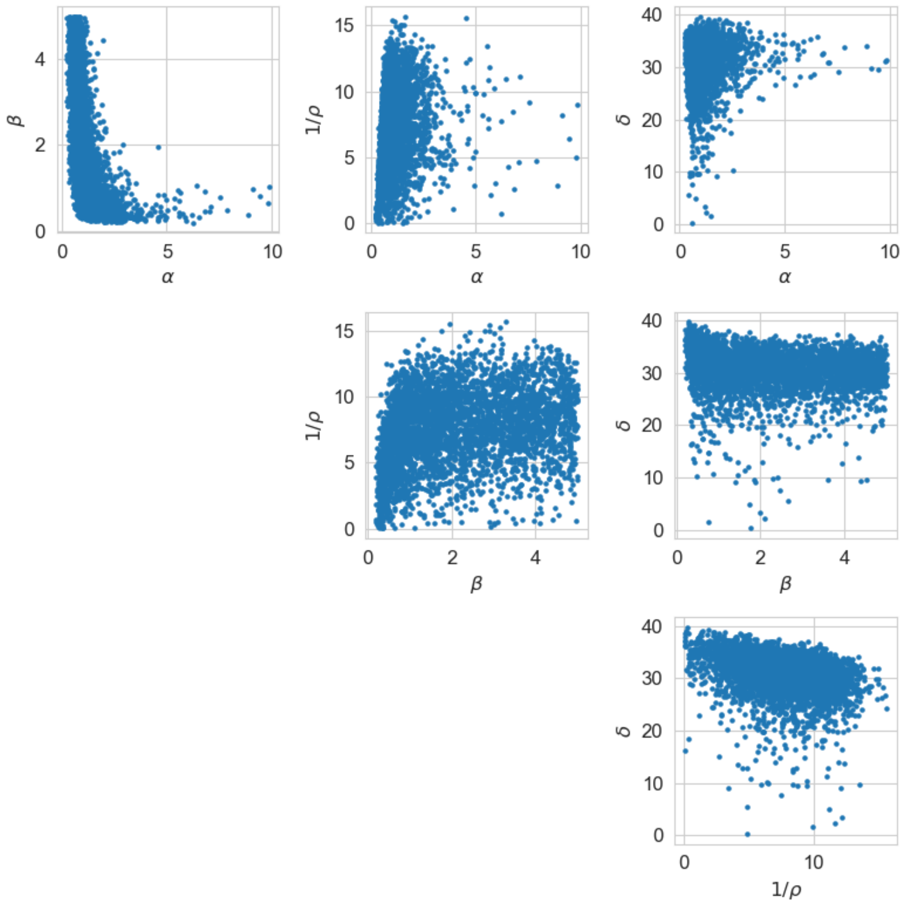


Figure 3: **S3: Correlations between the posterior parameters from the dispersal model.**

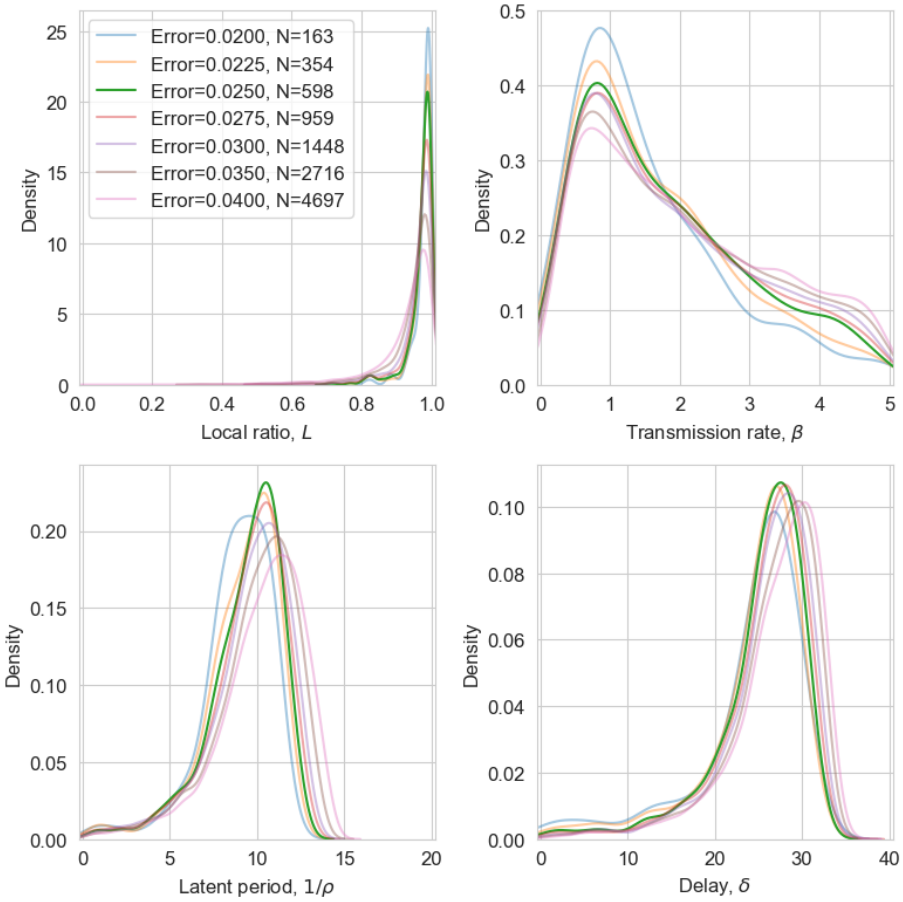


Figure 4: **S4: Posterior density plots for the lattice model for different error acceptance thresholds.** The default value of 0.025 is shown in bold, with a range of higher and lower values also presented. The distributions for each parameter appear quite consistent and, as is expected, there is a clear narrowing of the distributions as the threshold is reduced. The distributions appear to temporarily stabilise between values of 0.03 and 0.0225. While there is then some further movement for the lower threshold in some parameters, the number of acceptances here is rather low and we felt it a less reliable guide.

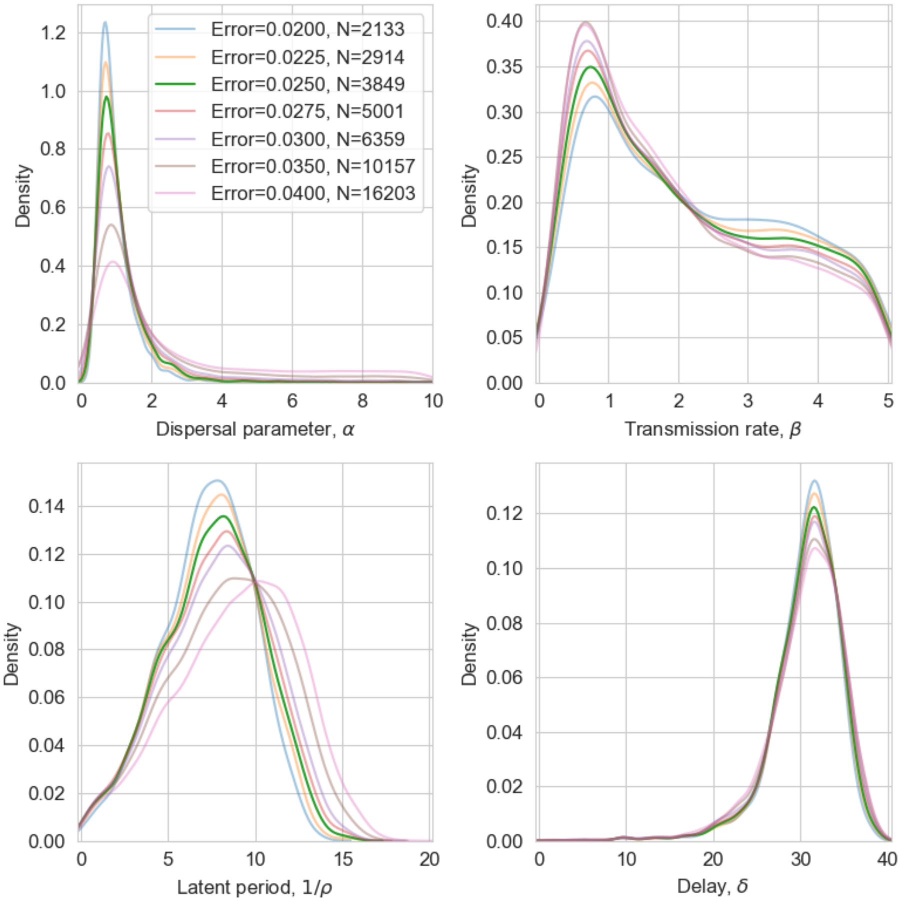


Figure 5: **S5: Posterior density plots for the dispersal model for different error acceptance thresholds.** The default value of 0.025 is shown in bold, with a range of higher and lower values also presented. The distributions for each parameter appear quite consistent and, as is expected, there is a clear narrowing of the distributions as the threshold is reduced. The distributions again stabilise between values of 0.03 and 0.0225.
